# Relationship between faecal microbiota and plasma metabolome in rats fed NK603 and MON810 GM maize from the GMO90+ study

**DOI:** 10.1101/593343

**Authors:** Robin Mesnage, Caroline I. Le Roy, Martina Biserni, Bernard Salles, Michael N. Antoniou

## Abstract

Safety concerns arising from the consumption of foods derived from genetically modified (GM) crops remains a highly debated and controversial subject. We report here a faecal microbiota compositional analysis in Wistar rats from the GMO90+ study, which fed glyphosate-tolerant NK603 (+/− Roundup application during cultivation) and Bt toxin MON810 GM maize for 6 months (at 11 and 33% composition of the feed) in comparison to their closest non-GM isogenic lines. We first integrated the faecal microbiota compositional data with results from plasma metabolomics to establish a baseline allowing us to understand which bacterial species can influence host metabolism. *Coriobacteriaceae* and *Acetatifactor* significantly predicted plasma metabolic profile in males, while *Bifidobacterium* and *Ruminococcus* were able to predict female plasma metabolites. We then investigated the differences in fecal microbiota composition between group of rats fed MON810 or NK603 GM maize varieties in comparison to their respective isogenic lines. Bacterial community richness was not altered by the test diets. There were no statistically significant differences in taxa abundance in the rat faecal microbiota that we could attribute to the consumption of either MON810 or NK603 GM maize varieties. In conclusion, we show that the consumption of the widely cultivated GM maize varieties NK603 and MON810 even up to 33% of the total diet had no effect on the status of the faecal microbiota compared to non-GM near isogenic lines.

## INTRODUCTION

Agricultural genetically modified (GM) crops are dominated by plants modified to tolerate herbicides and/or produce insecticides in order to facilitate pest management (Bonny, 2016). The majority of GM food crops currently being cultivated are maize and soybean varieties into which was introduced a modified version of the 5-enolpyruvylshikimate-3-phosphate synthase (EPSPS) from *Agrobacterium tumefaciens strain CP4* to confer tolerance to glyphosate-based herbicides. The EPSPS-CP4 transgene sidesteps the inhibition of the plant host equivalent enzyme, which is part of the shikimate pathway responsible for aromatic amino acid biosynthesis. An example of a glyphosate tolerant crop is NK603 maize, which is a GM event introduced in a large number of commercial varieties. The accumulation of glyphosate-based herbicide (GBH) residues by NK603 maize has been hypothesized to be a source of toxicity at the level of the gut microbiota. This is because some bacterial communities are equipped with the shikimate pathway and are thus sensitive to glyphosate (Lozano et al., 2018; Lu et al., 2013; Motta et al., 2018). However, the existence of negative effects from glyphosate exposure on the human microbiome, especially at typical, real-world levels of ingestion is still controversial and remains to be ascertained.

Insecticidal GM crops confer resistance to coleopteran insects and lepidopteran larvae by producing a mutated toxin modified from *Bacillus thuringiensis* species, collectively known as Bt toxins (Hilbeck and Otto, 2015). The MON810 event is used to confer insecticidal properties to maize by producing a modified cry1Ab delta-endotoxin. A number of laboratory animal feeding studies assessing potential toxicity of GM crops, including maize varieties, engineered with different Bt toxins conducted by academic groups (Finamore et al., 2008; Kilic and Akay, 2008) and by industry for regulatory purposes (de Vendomois et al., 2009), have reported changes in the structure and function of multiple organ systems. However, although a large number of studies have found that toxins derived from Bacillus thuringiensis cry proteins can have effects in mammals, as to whether these constitute either toxicity or can be deemed as innocuous remains to be resolved (Rubio-Infante and Moreno-Fierros, 2016). Concerns about the effect of Bt toxins on the gut microbiome remain since cry toxin fragments can be found in faeces from cows fed MON810-containing feed (Guertler et al., 2010). However, direct toxicity of cry toxins on mammalian microbiomes is unlikely since the toxin-specific receptor mediating the toxicity in insect epithelial cells is not detected in bacterial membranes (Nagamatsu et al., 1999; Oliveira et al., 2008).

Whether the consumption of foods derived from GM crops constitutes a health risk has been extensively investigated over the last 20 years and remains a highly debated and controversial subject (Domingo, 2016). Generally, there are three main possible sources of toxicity that can arise from GM foods. First, the foreign transgene product, such as Bt toxin insecticidal proteins, may directly cause harm. Second, increased residue levels and thus exposure to pesticides, particularly GBHs such as Roundup, to which the GM crop has been engineered to tolerate can give rise to ill-health. Third, altered plant biochemistry resulting from the imprecision of the GM transformation process and new combinations of gene function can lead to the unexpected production of novel toxins and allergens. Some reviewers have drawn attention to laboratory animal feeding studies that indicate signs of toxicity arising from diets containing GM crop ingredients (de Vendomois et al., 2009; Dona and Arvanitoyannis, 2009; Hilbeck et al., 2015; Krimsky, 2015), whilst other commentators report no safety issues connected GM foods (Delaney et al., 2018; Nicolia et al., 2014; Ricroch, 2013; Snell et al., 2012).

However, an area that remains relatively unknown are possible effects from the consumption of GM food crops on the gut microbial ecosystem (microbiome). All animal bodies are home to trillions of bacteria, fungi and viruses, having complex symbiotic or pathobiotic interactions with their hosts (Lloyd-Price et al., 2017). The gut microbiome is the largest of these communities, including approximately 3.8×10^13^ bacteria in humans (Sender et al., 2016). Alterations in human gut microbiome composition and function have been implicated in many diseases such as neurological, metabolic diseases as well as colorectal cancer (Clemente et al., 2012; Jackson et al., 2018; Murphy et al., 2019; Zitvogel et al., 2015). In addition, gut microbiome dysbiosis has been implicated in contributing to behavioural (e.g., autism, ADHD) and mental health (e.g., depression) problems (Dinan et al., 2015). Although the gut microbiome is partly under control of host genetics (Goodrich et al., 2014; Xie et al., 2016), its dynamics is mainly driven by environmental factors and in particular by diet (David et al., 2014; Rothschild et al., 2018; Zmora et al., 2019). The dietary intake in proteins, fat, carbohydrates, polyphenols and pre/probiotics have all been demonstrated to influence the gut microbiome with consequences on host immunologic and metabolic function (Singh et al., 2017). Well-known examples include anti-inflammatory effects of microbial-derived short chain fatty acids produced by the fermentation of polysaccharides (De Vadder et al., 2014). Some neuroactive compounds such as γ-aminobutyric acid are directly produced by certain gut bacteria (Dinan et al., 2015). Alternatively, gut bacteria are crucial regulators of serotonin production by enterochromaffin cells of the gut epithelium (Yano et al., 2015). Thus gut microbiome status can also impact the development of psychological disorders (Dinan et al., 2015; Osadchiy et al., 2019). Given the novel therapeutic possibilities offered by a modification of the gut microbiome through dietary interventions (Valdes et al., 2018; Zmora et al., 2019), it is essential to understand how the microbiome can change in response to different types of diets.

Some researchers have investigated the effects of a GM food-based diet on the composition of the faecal microbiota in laboratory rodents (Li et al., 2018) and non-human primates (Mao et al., 2016), as well as in farm animals such as cattle (Brusetti et al., 2011) and pigs (Buzoianu et al., 2012a; Buzoianu et al., 2012b). The most recent of these studies measured the composition of fecal microbiota in Sprague-Dawley rats fed two GM maize varieties that are representative of the two major GM crop traits (herbicide tolerance, herbicide tolerance + insecticide production), in comparison with their closest isogenic maize for 10 weeks (Li et al., 2018). There were no effects detected stemming from the consumption of these GM maize varieties on the overall health and diversity of fecal microbiota in comparison to their isogenic lines. Longer feeding trials may be necessary to assess life-long exposures to different diets although it has been noted that in general, approximately 70% of toxicological effects of chemical substances which are measured in two-year feeding trials can be predicted at 3-months from the start of exposure (EFSA, 2008).

Recently, results from a study, designated “GMO90+”, have been reported where laboratory rats were fed diets containing two different varieties of GM maize; MON810, which produces a type of Bt toxin and glyphosate-tolerant NK603 either sprayed or not sprayed with a Roundup GBH during cultivation (Coumoul et al., 2019). Wistar rats were fed diets containing NK603 (+/− GBH) and MON810 GM maize for 6 months at two concentrations (11 and 33% of the feed) and compared to diets containing the same quantities of the closest isogenic non-GM maize varieties. Measurements included in this study were transcriptomics (liver and kidney), metabolomics (plasma and urine), steroidomics (urine), as well as blood and urine biochemistry, necropsy, histopathology, and gut barrier integrity analysis. Overall, the most pronounced effect was attributed to gender differences followed by differences in the maize origin (MON810 grown in Spain compared to NK603 grown in Canada). Generally, pair-wise comparisons did identify some statistically significant differences between animals fed the GM maize compared to those consuming the non-GM closest isogenic lines. However, as no pattern of biological effects that could constitute potential signs of pathology were observed, and thus any long-term health implications stemming from these findings remains unknown.

In order to provide further insight into the effects of the GM maize diets in the GMO90+ study, we report here the faecal microbiota composition conducted by sequence analysis of 16S rRNA gene hypervariable regions, in the same animals used in this investigation (Coumoul et al., 2019). In addition, in order to understand the effects of the variations in gut microbiome composition on host health, we integrated the 16S rRNA gene amplicon sequencing data with the previously reported blood metabolomics data (Coumoul et al., 2019). Data from human cohorts indicate that the faecal metabolome largely reflects gut microbiome and its metabolic processes (Zierer et al., 2018). This approach can also help understand how differences in diet composition may have an effect on the host through the gut microbiome with potential pathological consequences.

We reveal sex-dependent relationships between the faecal microbiota and the plasma metabolome. However, none of these relationships were altered by ingestion of NK603 (+/−GBH) and the MON810 GM maize varieties.

## MATERIALS AND METHODS

### Rat feeding study

The animal feeding study was conducted at CiToxLAB (Evreux, France). The experimentation was approved by the French Ethical Committee (CETEA). The feces analysed in this study were obtained from Wistar Han RCC rats as previously described (Coumoul et al., 2019). Briefly, the animals were maintained at 22 ± 2°C under controlled humidity (50 ± 20%) and filtered, non-recycled air, with a 12 h-light/ dark cycle, with free access to food and water. Rats were kept in polycarbonate cages (Tecniplast 2154: polycarbonate with stainless steel lid; 940 cm2) containing autoclaved sawdust (SICSA, Alfortville, France). The experiment was performed blinded. The location of each cage within the experimental room was changed every week. After 14 days of acclimation with ACCLI diet (33% SyNepal), 6 weeks old Wistar Han RCC rats were fed a diet containing either 11% or 33% of GM NK603 (Pioneer 8906R) maize either treated or not treated with a glyphosate-based herbicide. Other test groups of animals were fed a diet containing either 11% and 33% of GM MON810 (DKC6667YG) maize. The GM diet groups were compared to animals fed an equivalent amount of near-isogenic control maize varieties (DKC6666 and Pioneer 8906 for MON810 and NK603 respectively). Details of maize production, as well as measurements of the presence of contaminants in the diets, and in the maize kernels, is detailed elsewhere (Chereau et al., 2018). Animals were randomized using a computerized stratification procedure, with mean body weights ± 10% between groups (per sex). This was changed weekly. The feces were collected by placing the animals in individual metabolism cages for 24 hours (at week 25/26).

### 16S MetaVx™ mammalian sequencing library preparation and Illumina MiSeq sequencing

The composition of the faecal microbiota was evaluated by measuring the faecal microbiota. A total of 189 fecal samples were available from rats at the 6-month termination point of the feeding period. The 16S MetaVx™ Mammalian next generation sequencing library preparations and Illumina MiSeq sequencing were conducted under contract with the commercial service provider GENEWIZ, Inc. (South Plainfield, NJ, USA). Fecal genomic DNA was extracted using a 3rd party kit. The resulting DNA samples were quantified using a Qubit 2.0 Fluorometer (Invitrogen, Carlsbad, CA) and DNA quality was checked by 0.6% agarose gel electrophoresis. Sequencing libraries was prepared using the 16S MetaVx™ Mammalian Library Preparation kit (GENEWIZ, Inc., South Plainfield, NJ, USA). Briefly, 50 ng DNA was used to generate amplicons that cover V3 and V4 16S rDNA gene hypervariable regions of bacteria and Archaea. Indexed adapters were added to the ends of the 16S rDNA amplicons by limited cycle PCR. Sequencing libraries were validated using an Agilent TapeStation (Agilent Technologies, Palo Alto, CA, USA), and quantified by Qubit and real time PCR (Applied Biosystems, Carlsbad, CA, USA). DNA libraries were multiplexed and loaded on an Illumina MiSeq instrument according to the manufacturer’s instructions (Illumina, San Diego, CA, USA). Sequencing was performed using a 2×250 paired-end (PE) configuration; image analysis and base calling were conducted by the MiSeq Control Software (MCS) on the MiSeq instrument. Three technical replicates were performed and a total of 164,205 +/− 52,440 reads were obtained for each sample.

### Plasma metabolomics

The plasma metabolomics has been described in a preceding publication (Coumoul et al., 2019) and was performed under contract with Profilomics (Huningue, France). Briefly, a liquid chromatography–high resolution mass spectrometry (LC–HRMS) approach was used. The plasma samples were separated on a HTC PAL-system coupled with a Transcend 1250 LC instrument using a Sequant ZICpHILIC column. The column effluent was injected into the heated electrospray source of a Q-Exactive mass spectrometer.

### Processing of 16S rDNA gene sequencing data

The DADA2 algorithm (version 1.6) was first used to correct for sequencing errors and derive exact sequence variants (ESV) using the R version 3.4.1. ESV are a higher-resolution version of the operational taxonomic unit (OTU). While OTUs are clusters of sequences with 97% similarity, ESVs are unique sequences (100% similarity). The use of ESV is being advocated to replace OTU in order to improve reproducibility between studies because a threshold of 97% does not allow a reproducible classification of closely related bacteria (Callahan et al., 2017).

Each of the three replicates were first processed separately in order to evaluate the differences between each Miseq run. Since the quality of these three sequencing runs was very similar with no batch effects, the 3 technical replicates were merged to reduce variability by effectively averaging over the technical replicates. Forward and reverse reads were trimmed by 5 and 10 bp, respectively. Error rate convergence values reached 0.007 and 0.008 for forward and reverse reads, respectively. A total of 91% of the reads successfully merged. We filtered out sequence variants with more than 95% missing values. The taxonomy was assigned using the SiLVA ribosomal RNA gene database v128. We performed a multiple-alignment using the *DECIPHER* R package in order to create a phylogenetic tree (Figure 1A) with the *phangorn* R by fitting a GTR+G+I (Generalized time-reversible with Gamma rate variation) model on a neighbour-joining tree. The count table, the metadata, the sequence taxonomies, and the phylogenetic tree were ultimately combined into a single R object, which was analysed using the *phyloseq* package (McMurdie and Holmes, 2013). These data have been submitted to the BioSample database and are accessible through accession number PRJNA528035.

**Figure 1.**
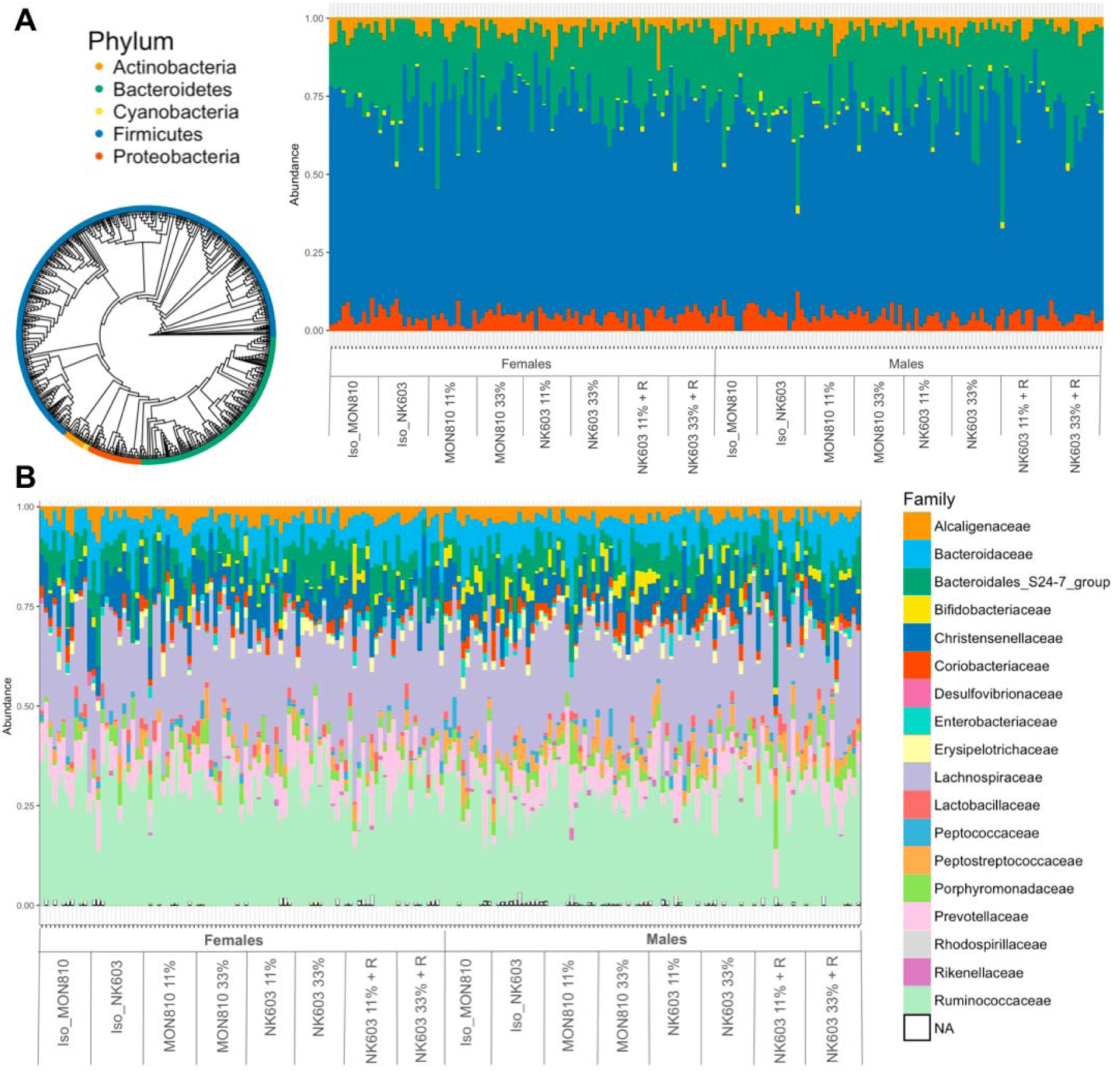
Taxonomic composition of the faecal microbiome of rats fed GM maize varieties MON810 and NK603. Feces from rats fed for 6-months GM maize MON810 and NK603. (cultivated with or without Roundup application, NK603 + R) at two doses (11 and 33%), in comparison to their closest isogenic (Iso_MON810, Iso_NK603), were analysed using 16S rRNA gene sequencing. **A**. Phylogenetic tree showing a good clustering of the different phyla. **B**. Analysis at the bacterial family level reveals a good taxonomy assignment where only few ESV remain unidentified (NA). No differences between the different test and control groups is evident at this level.

### Statistical analysis

The statistical analysis was performed in R version 3.4.1. We have followed the Bioconductor workflow developed by Benjamin Callahan (Callahan et al., 2016). The alpha diversity was measured using the Shannon index calculated with the plot_richness *phyloseq* function, with statistical significance assessed with a Kruskal-Wallis rank sum test. The beta diversity was visualized by a MultiDimensional Scaling (MDS) ordination of the sample Bray-Curtis distances. We investigated the statistical significance of differences in taxa abundance by fitting an ANOVA model using the R *lm* function, with the sex of the animal as covariate.

We investigated the prediction power of the 72 genera on the faecal metabolome. Metabolomics data were mean centred and normalised prior to analysis. Prior to supervised analysis we conducted an unsupervised principal component analysis (PCA) to identify outliers and potential covariates. We observed that male and female rats had very distinct metabolic profiles as previously reported on this dataset (Coumoul et al., 2019). We therefore pursued the analysis separately for male and female animals. Prediction potential of each taxa was assessed individually by O-PLS DA (Wold et al., 2004) using the metabolomics profiles as a matrix of independent variables and genera as predictors. Models presenting a goodness of prediction coefficient (Q^2^Y) above zero were selected for validation using a thousand permutation. Models were considered as significant if the permutation returned P < 0.05.

## RESULTS

The fecal samples analysed in this study were obtained from rats that formed a 6-month feeding trial looking at physiological effects from the consumption of MON810 and NK603 GM maize varieties by comparison with diets containing equivalent amounts of near isogenic non-GM maize (Coumoul et al., 2019).

The processing of the Illumina MiSeq PE250 16S rRNA gene sequencing data provided an average of 41,085 +/− 10,352 clean reads per sample, with a mean length of 427 bp. A total of 1530 ESV were detected. These corresponded to 1529 bacterial ESV and 1 archea (Methanobacteria). A total of 6 different phyla were detected (Figure 1A). The faecal microbiota of the rats analysed in this study was dominated by Firmicutes, more specifically by Ruminococcaceae and Lachnospiraceae families (Figure 1B). Overall, a total of 72 and 19 distinct genus and species were identified.

We began by integrating the composition of the faecal microbiota with the blood plasma metabolomics data obtained from the same animals (Coumoul et al., 2019) in order to understand which taxa influenced the composition of the metabolome. This is an important baseline to assess the biological relevance of any statistically significant differences detected in the composition of the faecal microbiota. Using an unsupervised multivariate analysis showed a clear separation between male and female metabolic profiles (Figure 2). The loading of PC1 (Supplementary Table 1) revealed that the metabolic gender separation was mostly driven by increased carnitine, methylhistamine, butyril-carnitine, L-threonic acid, carnosine and acetylcholine in males and an increase in lysine, 5-methyldeoxycytidine, corticosterone glycyl-L-leucine, N6-acetly-L-lysine, tryptophane, perillic acid and indoleacrylic acid in female animals. Separating males from females we identified 2 genera in males (Coriobacteriaceae and *Acetatifactor*) and 2 in females (*Bifidobacterium* and *Ruminococcus)* able to significantly predict plasma metabolic profiles. Coriobacteriaceae relative abundance was significantly associated with 29 metabolites (Figure 3A and 3B) including L-anserine, carnosine, leucine/proline (positively) and allantoin cinnamic acid perillic acid (negatively). *Acetatifactor* was significantly associated with 30 metabolites including one bile acid (lithocholic acid-3 sulphate –positive-) and three N-acetyl amino acids (arginine, asparagine and glutamine –positive-). In female rats, *Bifidobacterium* was positively associated with four bile acids (deoxycholic acid, cholic acid, lithocholic acid and lithocholic acid 3 sulphate). Finally, in female animals, *Ruminococcus* was the genera predicting the widest proportion of the plasma metabolome with 39 metabolites significantly associated with its relative abundance.

**Figure 2.**
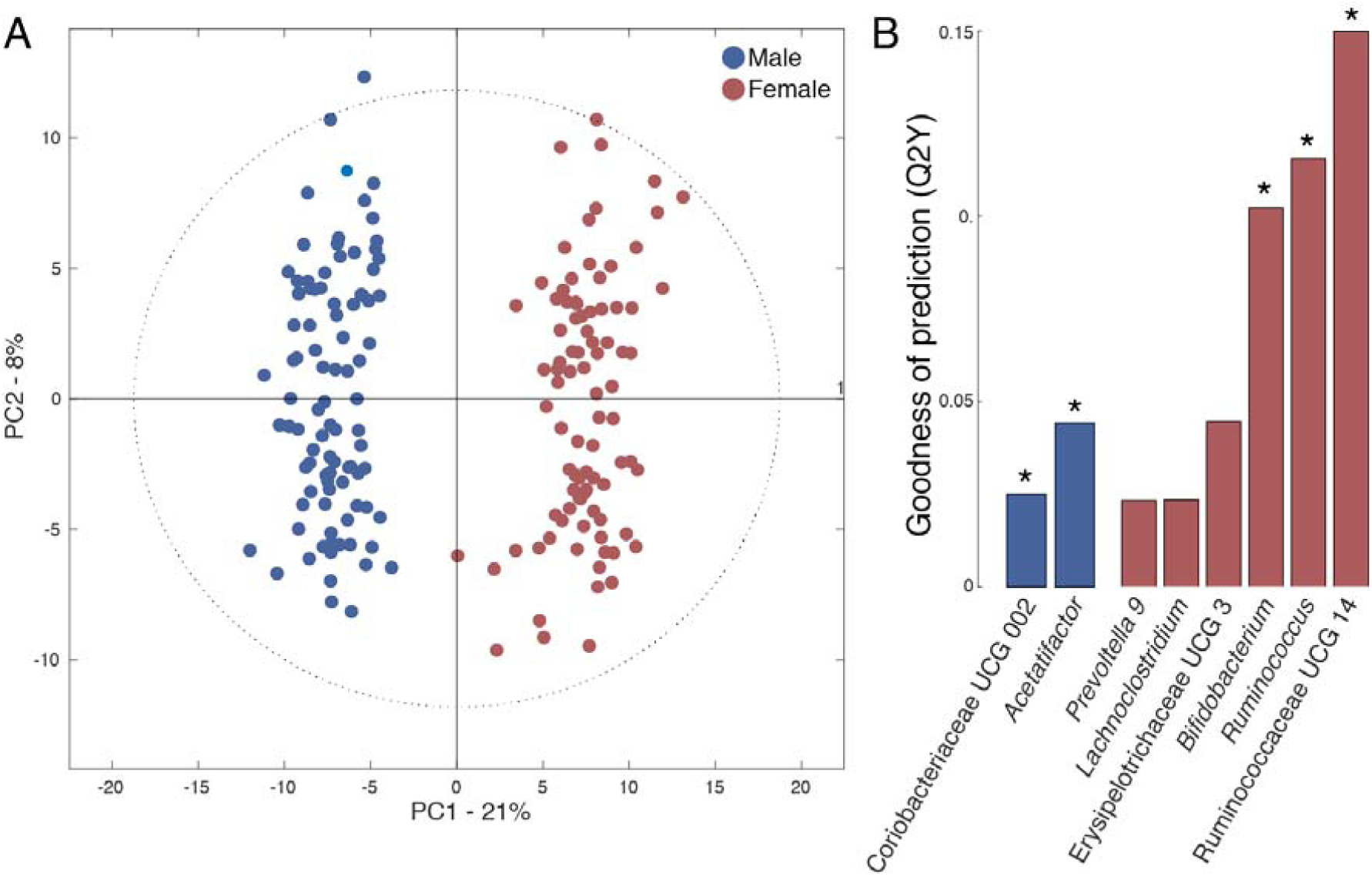
Gender variation in faecal bacteria-blood metabolome prediction. A. PCA score plot on PC1 and PC2 of plasma metabolome composition. B. Goodness of prediction of the O-PLS DA models using genera as predictors and plasma metabolome profiles as a matrix of independent variables in male (blue) and female (pink) animals separately. * P < 0.05 after permutation.

**Figure 3.**
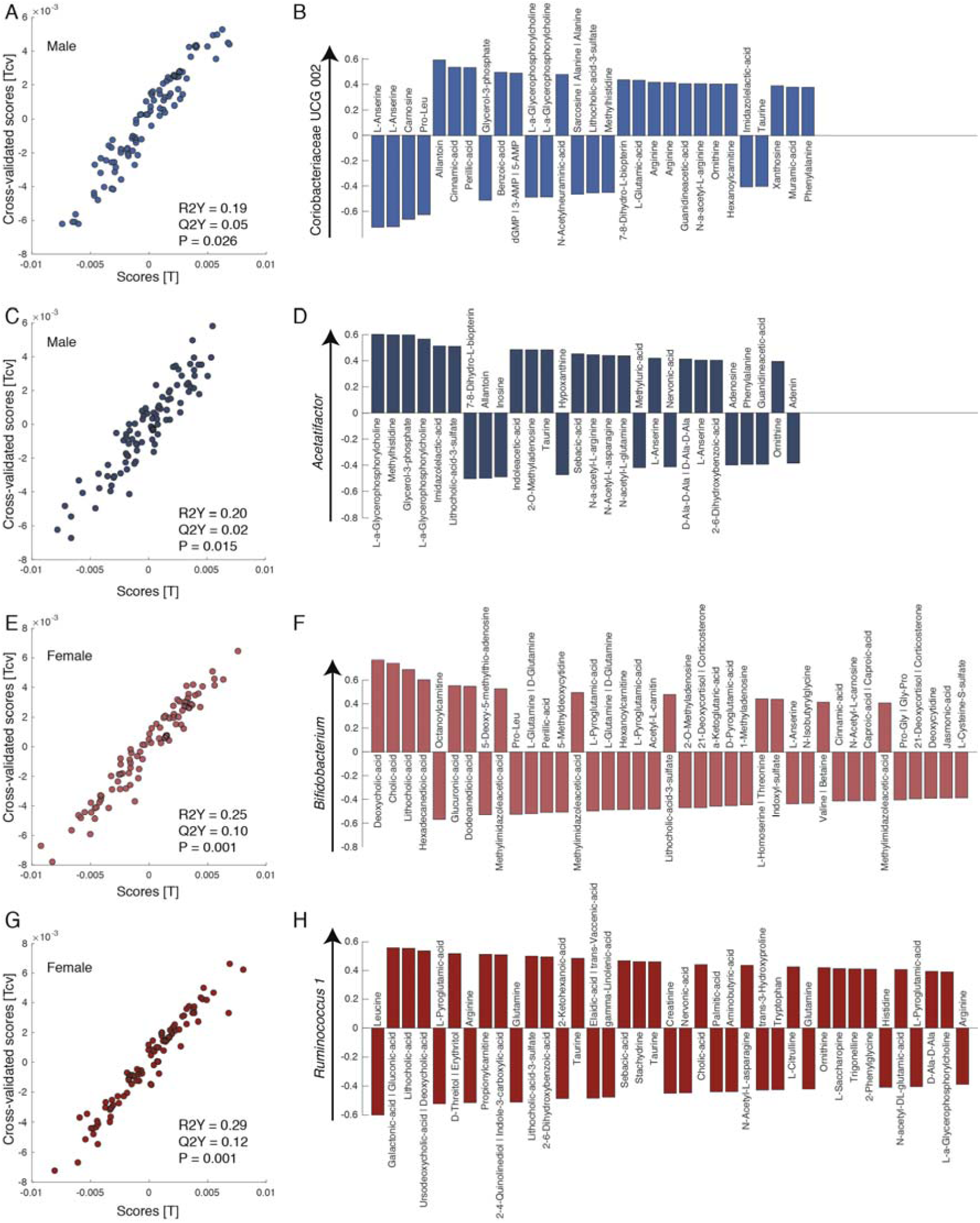
Faecal bacteria are predictors of plasma metabolite profile. A. Scores vs. cross validate scores of the O-PLS DA model generated using Coriobacteriaceae UCG 002 relative abundance as a predictor of plasma metabolomics profile in male mice. B. Bar plot displaying the metabolites significantly (P < 0.05/279 –Bonferroni-) contributing to the O-PLS DA model described in A. Metabolites are ordered by level of significance. Metabolites pointing up are positively associated with Coriobacteriaceae relative abundance while metabolites pointing down are negatively associated with Coriobacteriaceae relative abundance. C, E and G were obtained as described for A using *Acetatifactor, Bifidobacterium* and *Ruminococcus* 1 as predictor respectively. D, H and F were obtained as described in B.

Bacterial community richness in the faecal microbiota of rats fed diets with GM and non-GM maize varieties was studied using the Shannon diversity index (Figure 4). There was no significant difference between the different groups after a statistical evaluation using a Kruskal-Wallis rank sum test. However, females had a lower alpha diversity compared to males (p = 0.032). We further performed an analysis of Bray-Curtis distances between the samples (Figure 5). First, this analysis indicated that there were no outliers. Second, the sample plot shows a clustering of the treatment group indicating that a 6-month feeding period with the different maize diets had not caused changes in microbiota composition. The gender difference becomes visible at this level, with the male samples having a higher weight on the first component compared to females.

**Figure 4.**
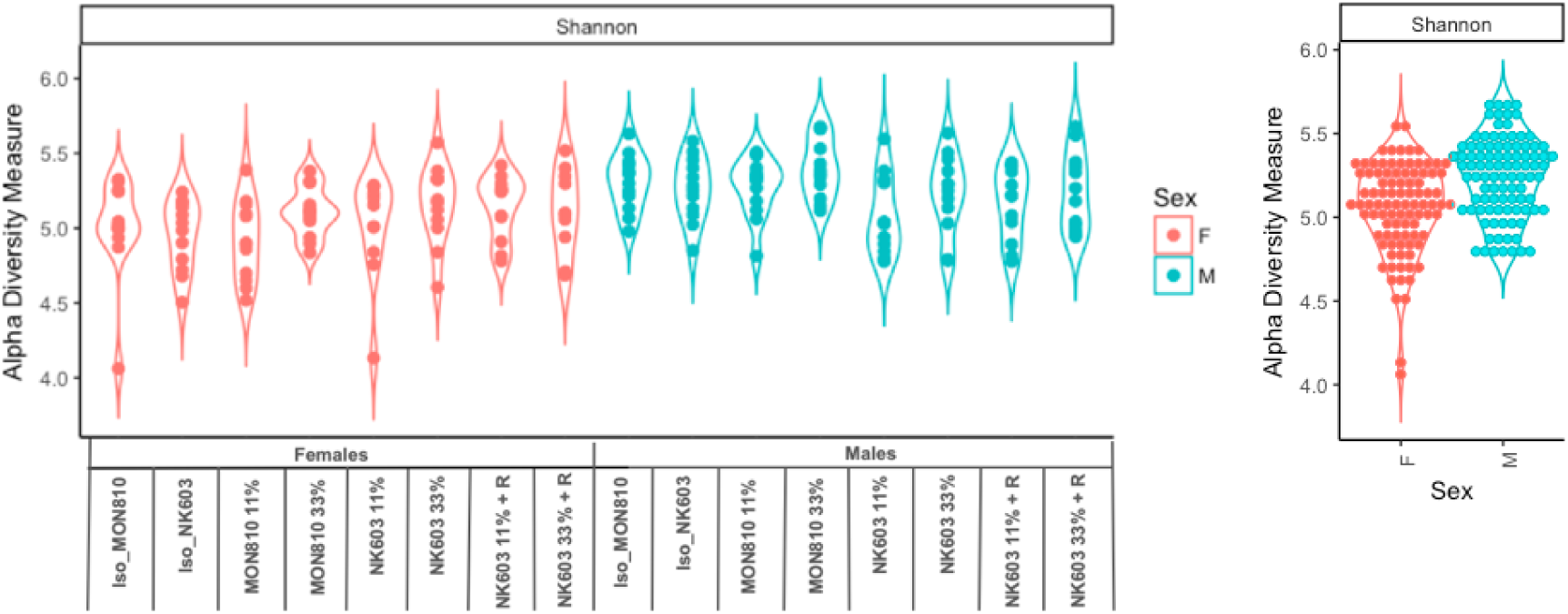
Bacterial community richness in the faecal microbiota of rats fed GM and non-GM maize. Rats were fed a diet containing either 11% or 33% of GM maize varieties MON810 or NK603 either with (NKG) or without (NK) Roundup application and compared to animals consuming an equivalent amount of nearest isogenic non-GM maize. Faecal microbiota composition was determined by 16S rRNA gene sequencing of fecal samples collected at the 6-month termination time point of the study. Richness was calculated using the Shannon diversity index in Phyloseq.

**Figure 5.**
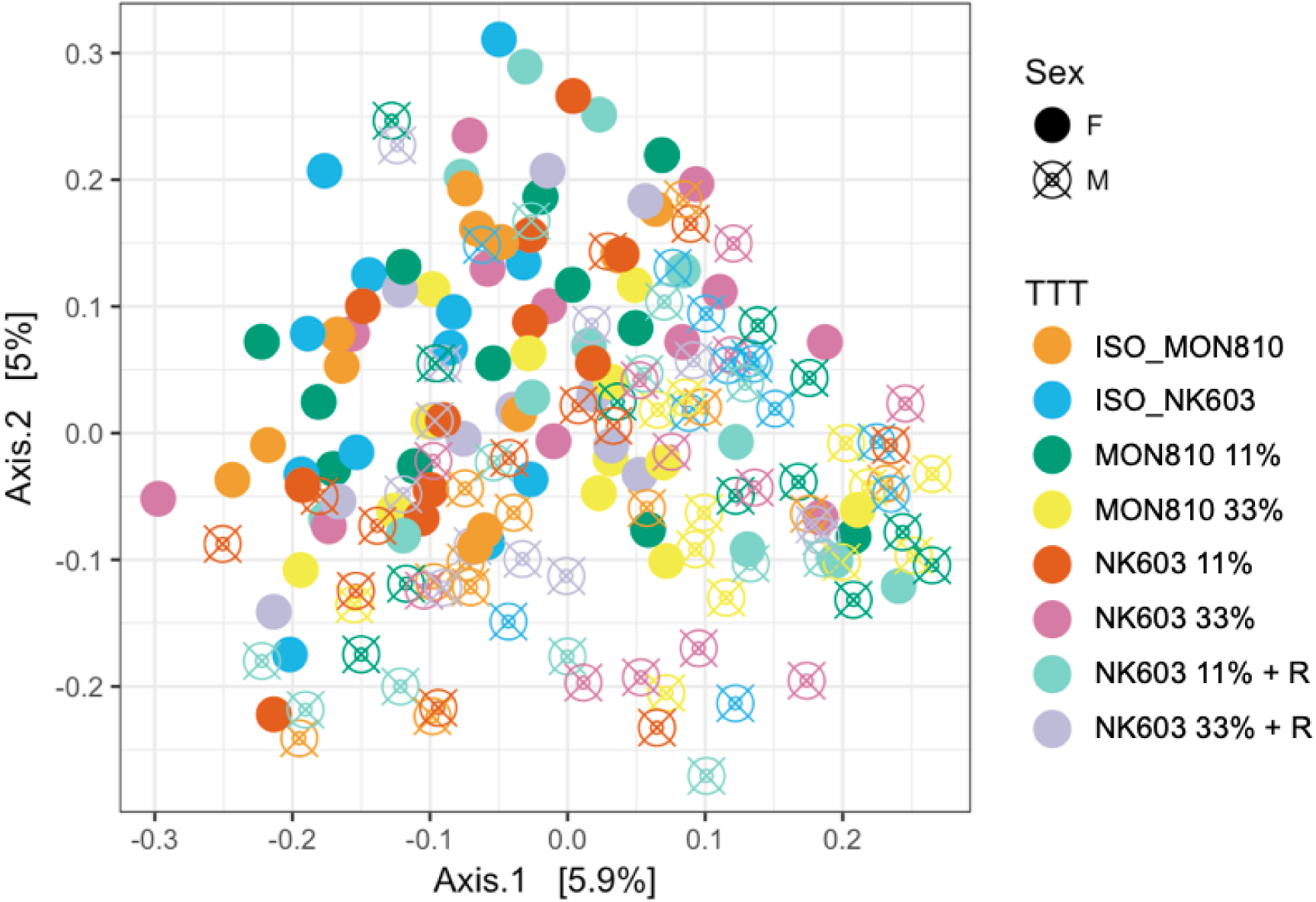
Comparison of microbial community composition between faecal samples from rats fed GM and non-GM maize. Rats fed a diet containing either 11% or 33% of GM maize varieties MON810 or NK603 either with (NKG) or without (NK) Roundup application were compared to animals consuming an equivalent amount of nearest isogenic non-GM maize. Faecal microbiota composition was determined by 16S rRNA gene sequencing of faecal samples collected at the 6-month termination time point of the study. Multi-dimensional scaling ordination of Bray-Curtis distances. Although the treatment groups do not cluster together, sex-specific differences in faecal microbiota composition was evident.

We also investigated the effects of the MON810 and NK603 GM maize-containing diets at the level of taxa by undertaking a multivariate analysis of differential taxa abundance. There were no statistically significant differences in taxa abundance that we could attribute to the effects of the MON810 (Supplementary Material 2 and 3) and the NK603 (Supplementary Material 4 and 5). Thus, the consumption of MON810 or NK603 did not change the composition of the faecal microbiota.

## DISCUSSION

The question of the safety of GM crop consumption is a burning question with large societal, economic and political consequences for agricultural and food systems. Whether there are differences in health effects between GM crop consumption and their isogenic lines is a controversial issue with no agreed scientific consensus on this (Hilbeck et al., 2015; Krimsky, 2015; Nicolia et al., 2014; Ricroch, 2013).

A potential effect of GM food on blood metabolome was not observed, which in turn could have modified the gut microbiome taxonomic richness and evenness (Coumoul et al., 2019). However, here we address the possibility of a direct effect of GM food on the gut microbiome, which could potentially lead to negative health outcomes. We present the results of the longest (6 month) investigation of the effects from consuming glyphosate-tolerant NK603 and Bt toxin MON810 GM maize crops on the composition of the faecal microbiota in Wistar Han RCC rats. No alterations in microbiota composition were detected stemming from the consumption of these GM crops in comparison to diets with equivalent amounts of their respective non-GM isogenic maize varieties. This is similar to findings of a previous study, which also reported no changes in the rat microbiome from the consumption of different GM maize varieties (Li et al., 2018).

We integrated the results we obtained from the faecal microbiota analysis and blood metabolome data from the same rats (Coumoul et al., 2019) and found that a number of bacterial taxa could be used as predictors of the animal’s plasma metabolite profile. Interestingly, we observed that faecal microbes affected host systemic metabolism in a gender specific manner. Indeed, bacterial predictors were gender specific and affected distinct metabolites. The metabolic profile in males was dominated by markers of protein metabolism (carnitine, carnosine and butyryl-carnitine) that may reflect their higher muscular mass compared to females. Additionally, we observed that only in males, Coriobacteriaceae relative abundance was positively associated with allantoine. Both Coriobacteriaceae and allantoine increase following exercise (Batacan et al., 2017; Hellsten et al., 1997; Zhao et al., 2018). In females, we observed that *Bifidobacterium* was positively associated with several primary and secondary bile acids as was *Ruminococcus*. Primary bile acids are produced by the liver from cholesterol and excreted into the small intestine. They are then transformed into secondary bile acids by gut bacteria and reabsorbed by the host. *Bifidobacterium* is known for its cholesterol lowering properties (Bubnov et al., 2017), which may explain the observed associations. Although it is not directly relevant for the effects of GM corn, this set of data can inform on general aspects of the interface between the gut microbiome and the blood metabolome in laboratory rodents.

The rat fecal samples analysed in this investigation were obtained from, and thus add to, the results obtained from the GMO90+ research programme (Coumoul et al., 2019). The GMO90+ study found a number of statistically significant differences in various physiological parameters and expression of some genes in the GM maize fed groups compared to control animals. However, no clear effects on distinct biological systems that could signify negative health outcomes were seen. Thus, if there any health implications stemming from these statistically significant differences, especially in the long term, remains unknown. A previous study has reported potentially negative health effects from feeding Sprague-Dawley rats a variety of NK603 maize (+/− Roundup application) over a 2-year period (Seralini et al., 2014). A follow-up investigation (designated as G-TwYST) also of 2 years duration, rats were fed the same cultivation of NK603 GM maize as that used in the GMO90+ study (Steinberg et al., 2019). No consistent differences were observed between test and control groups in this G-TwYST study that were indicative of negative health effects (Steinberg et al., 2019). However, as the GMO90+ and G-TwYST studies and the investigation by Séralini and colleagues possess some major differences in design (e.g., strain of rat used, number of rats per diet group, study duration) they constitute distinct studies, with a direct comparison of reported outcomes between them difficult.

The NK603 GM corn used in this investigation was cultivated with or without the application of a Roundup GBH at the pre-and post-emergence stages of the cropping cycle (Coumoul et al., 2019). A number of studies have investigated the effects of glyphosate or its commercial herbicide formulations on the rat faecal microbiota with some showing changes in the microbiota composition (Lozano et al., 2018; Mao et al., 2018; Nielsen et al., 2018). In the study presented here, glyphosate was detected in all maize feeds regardless of whether the grain had been sprayed or not sprayed with Roundup GBH during cultivation (Chereau et al., 2018). As only the NK603 maize sprayed with Roundup was found to contain low levels (16μg/kg) of glyphosate (Chereau et al., 2018), this implies that the generally higher glyphosate contamination of the feeds (50-75μg/kg) is coming from other standard components (e.g., cereals, soybeans) of the feed formulation as previous detected (Mesnage et al., 2015). Consequently, this experiment cannot be used to draw conclusions on effects of GBH residues on the gut microbiome. Further experiments will thus be needed to ascertain if glyphosate or its commercial formulations such as Roundup can cause changes in the gut microbiome.

The study of gut microbial ecology in human health and diseases is a rapidly growing field of research receiving attention from researchers and the general public. It is a very young discipline without consensus “best practices” and with methodologies which are constantly evolving (Pollock et al., 2018). We evaluated the composition of the gut microbiome thought an evaluation of the faecal microbiota. However, gut microbiome metabolism cannot be fully reflected by stool composition (Zmora et al., 2018). The use of additional omics methods such as metatranscriptomics and metabolomics could provide more insights into gut functional alterations by bringing the biological significance of molecular profiles closer to phenotype. This can be the case when some bacteria are highly abundant but they are transcriptionally inactive, whereas some are very active despite their low abundance (Schirmer et al., 2018).

In conclusion, our study in a mammalian, rat model system with animals fed a 33% maize based diet, showed that faecal microbiota composition was closely correlated with the blood metabolite profile in a gender-specific manner with *Coriobacteriaceae* and *Acetatifactor* significantly being able to predict the plasma metabolic profile in males, while *Bifidobacterium* and *Ruminococcus* were able to predict the spectrum of plasma metabolites in females. We did not observe any difference in the fecal microbiota composition between control non-GM maize fed groups and GM maize test groups. In addition, no effect on the fecal microbiota was seen when comparing the GM Bt toxin MON810 and Roundup tolerant NK603 maize groups, even though these contained different contaminants and slightly different metabolome signatures (Bernillon et al., 2018). In summary, we found that the consumption of the widely used GM maize events NK603 and MON810 even up to 33% of the total diet had no effect on the status of the fecal microbiota composition compared to non-GM near isogenic lines.

## Acknowledgments

This work was funded by the Sustainable Food Alliance (USA) whose support is gratefully acknowledged.

## SUPPLEMENTARY MATERIAL

**Supplementary Material 1. Loading of the Principal Component 1 showing a metabolic gender separation.**

**Supplementary Material 2. Statistical significance of the changes in relative abundance for bacterial genera after the administration of the MON810 maize in rats.**

**Supplementary Material 3. Changes in relative abundance for bacterial genera after the administration of the MON810 maize in rats.** We presented the statistical comparisons with the lowest p-value (Supplementary Material 2). There was no statistical difference in faecal microbiota composition can be attributed to the inclusion of MON810 in rat diet.

**Supplementary Material 4. Statistical significance of the changes in relative abundance for bacterial genera after the administration of the NK603 maize in rats.**

**Supplementary Material 5. Changes in relative abundance for bacterial genera after the administration of the NK603 maize in rats.** We presented the statistical comparisons with the lowest p-value (Supplementary Material 4). There was no statistical difference in faecal microbiota composition can be attributed to the inclusion of MON810 in rat diet.

