## Supplementary figures and images for "Relationship between faecal microbiota and plasma metabolome in rats fed NK603 and MON810 GM maize from the GMO90+ study"

### Supplementary Material 3

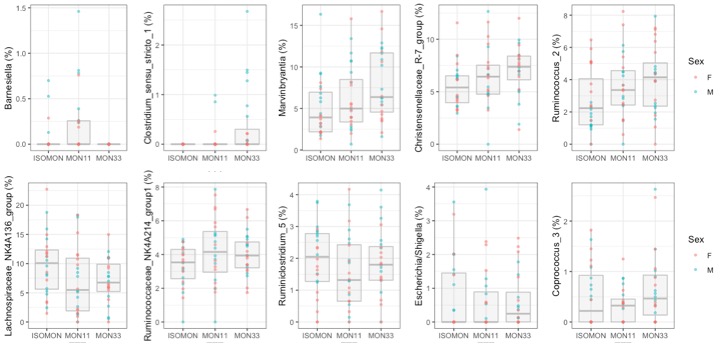

### Supplementary Material 5

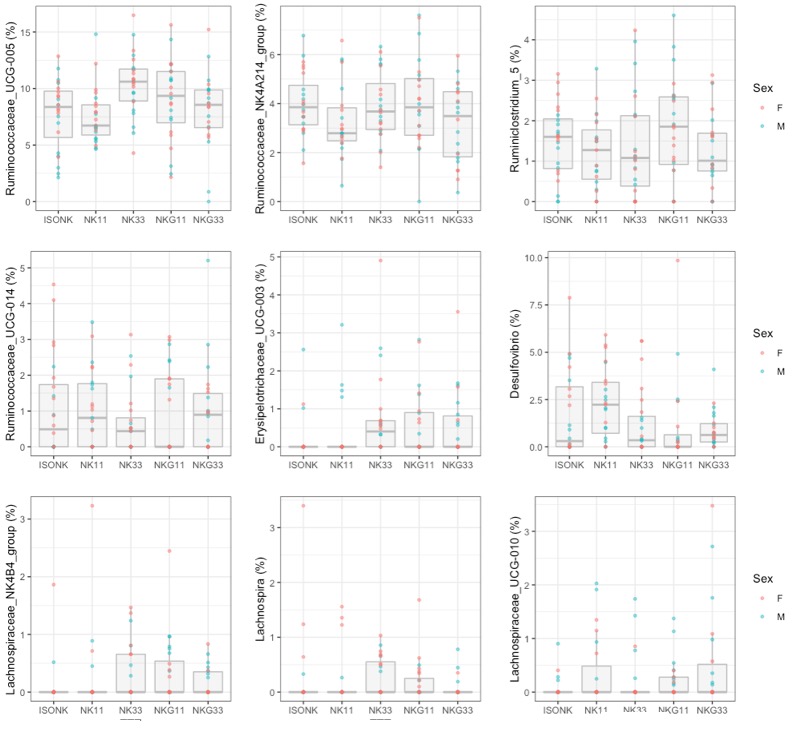
